# Organic matter influence on ooid formation: New insights into classic examples (Great Salt Lake, USA; Triassic Germanic Basin, Germany)

**DOI:** 10.1101/2023.04.11.536455

**Authors:** Yu Pei, Pablo Suarez-Gonzalez, Jan-Peter Duda, Joachim Reitner

## Abstract

Ooids are particles composed of a tangential or radial cortex growing around a nucleus. They are common in carbonate deposits of almost any geological age and provide insights into environmental conditions. However, abiotic or biotic factors influencing their formation remain unclear. This study aims to advance our understanding of ooid formation with a multi-analytical approach (e.g., FE-SEM, Raman spectroscopy, μ-XRF) to classic examples from Great Salt Lake (GSL; USA) and the Lower Triassic Germanic Basin (GB; Germany). Both deposits represent hypersaline shallow-water environments where ooids are closely associated with microbial mats. GSL ooids are dominantly 0.2–1 mm in size, ellipsoidal to subspherical in shape, composed of aragonite, and contain organic matter (OM). GB ooids are mainly ≤4 mm in size, spherical to subspherical in shape, composed of calcite, and currently contain little OM. Despite the differences, both ooids have the same cortex structures, likely reflecting similar formation processes. Some GSL ooids formed around detrital grains while others exhibit micritic particles in their nuclei. In GB ooids, detrital nuclei are rare, despite the abundance of siliciclastic particles of various sizes in the host rocks. GB deposits also include “compound ooids”, i.e., adjacent ooids that coalesced with each other during growth, suggesting static *in-situ* development, which is supported by the lack of detrital grains as nuclei. GB ooids also grew into laminated microbial crusts with identical microstructures, further indicating a static formation. Such microbial crusts typically form through mineral precipitation associated with OM (e.g., extracellular polymeric substances), suggesting a similar formation pathway for ooids. The inferred key-role of OM is further supported by features in radial ooids from the GSL, which commonly exhibit, from their nuclei towards their surface, increasing OM contents and decreasing calcification.

## 1 Introduction

Ooids are significant particles of carbonate deposits from many different sedimentary environments since the Archaean (e.g., Kalkowsky, 1908; Davies et al., 1978; Simone, 1981; Krumbein, 1983; Summons et al., 2013; O’Reilly et al., 2017; Siahi et al., 2017; Mariotti et al., 2018; Diaz & Eberli, 2019; Flannery et al., 2019). They are typically characterized by tangential and/or radial cortices and have sizes of < 2 mm (Richter, 1983; Flügel, 2010). In some cases, ooids can reach sizes of > 2mm (“giant ooids”: e.g., Sumner & Grotzinger, 1993; Li et al., 2015; 2021). Ooids have caught the attention of humans since prehistoric times (Binsteiner et al., 2008; Weber et al., 2022) and were already mentioned by Roman naturalists (Burne et al., 2012). However, their first scientific description was provided in a treatise devoted entirely to the ooids of the Lower Buntsandstein (Lower Triassic) from northern and central Germany by Brückmann (1721). He followed Volkmann (1720) in using the Greek word “oolithos” (literally “egg-rock”, due to their similarity to fish roe), a translation of the German “Rogenstein” or “Eierstein” (Burne et al., 2012). In his landmark book on the microfacies of carbonate rocks, Flügel (2010) defined ooids as “spherical and egg-shaped carbonate or non-carbonate coated grains exhibiting a nucleus surrounded by an external cortex, the outer part of which is concentrically smoothly laminated”.

Despite this long history of research, the formation of ooids still remains disputed and unresolved, and both inorganic and organic hypotheses have been proposed. The inorganic hypothesis involves a direct precipitation of aragonite or calcite from fluids supersaturated with respect to CaCO3 (e.g., Illing, 1954; Sumner & Grotzinger, 1993; Duguid et al., 2010; Trower et al., 2017, 2018). In this model, larger ooids are predicted to result from faster precipitation rates promoted by a higher calcium carbonate saturation state, combined with increased agitation to allow larger grains to be transported for active growth (Trower et al., 2017; Li et al., 2021). In the organic hypothesis, aragonite or calcite precipitation is closely associated with microbial extracellular polymeric substances (EPS), which consist of complex mixtures of various organic compounds such as polysaccharides, proteins, nucleic acids and lipids (Flemming & Wingender, 2010; Flemming, 2016; Decho & Gutierrez, 2017). EPS is widespread in biofilms (Wingender et al., 1999; Flemming et al., 2007; Neu & Lawrence 2010), implying the presence of a rich and diverse community of microorganisms during ooid formation (Friedman et al., 1973; Gerdes et al., 1994; Brehm et al., 2006; Plée et al., 2008; Summons et al., 2013; Woods, 2013; Diaz et al., 2015; 2017; O’Reilly et al., 2017; Hubert et al., 2018). Microorganisms are put forward to induce carbonate precipitation by their metabolic activity. However, living microorganisms must not necessarily be directly involved, as carbonate precipitation can as well be linked to degraded organic matter (OM) (“Organomineralization”: Mitterer, 1968; Suess & Fütterer, 1972; Ferguson et al., 1978; Trichet & Défarge, 1995; Reitner et al., 1995a, b, 1997; Reitner, 2004).

Herein, we study Lower Triassic ooids from the Germanic Basin (GB; Germany) and compare them with modern counterparts from the Great Salt Lake (GSL; Utah, USA). Ooids from both localities have been continuously studied since the XIX century until now (e.g., Rothpletz, 1892; Kalkowsky, 1908; Kässbohrer & Kuss, 2019; 2021; Ingalls et al., 2020, Trower et al., 2020), and have been central for the development of early hypotheses about the organic influence on ooid formation (e.g., Kalkowsky, 1908 for GB ooids; Rothpletz, 1892 for GSL ooids). Although more than a century has passed since those early hypotheses, much remains unknown about the exact role of OM in forming ooids. This work aims at advancing that direction by analyzing the similarities and differences between ooids from the very same classic localities that originated the idea of organic influence on ooids, and by discussing their possible formation processes.

## 2 Materials and methods

### 2.1 Geological setting and sample material

The Early Triassic Buntsandstein Group was deposited in the GB, which extended from England to Belarus and from Denmark to Germany (Figure 1a,b). It was mainly a continental basin but occasionally connected to the western Tethys Ocean (Meliata Ocean), and characterized by subtropical to arid climates (Ziegler, 1990; Stampfli, 2000; Weidlich, 2007; Scholze et al., 2017; Scotese, 2021). This study focuses on the Lower Buntsandstein Subgroup in Germany (Fig. 1b). The subgroup can be up to ∼450 m thick, consisting of siliciclastic rocks intercalated with carbonates (Ziegler, 1990). It is further subdivided into the Calvörde and Bernburg Formations. The studied ooids and stromatolites occur in the Bernburg Formation. They may be formed in a lacustrine environment characterized by elevated salinities (Paul & Peryt, 2000; Käsbohrer & Kuss, 2019), although some studies favour a marine environment due to a transgression (Weidlich, 2007). In addition to materials sampled during various field campaings between 2018 and 2020 at the Heeseberg and surroundings, and in Beesenlaubingen (Figure 1b), we studied existing samples that are archived in the Göttingen Geoscience Collections at the University of Göttingen.

**Figure 1.**
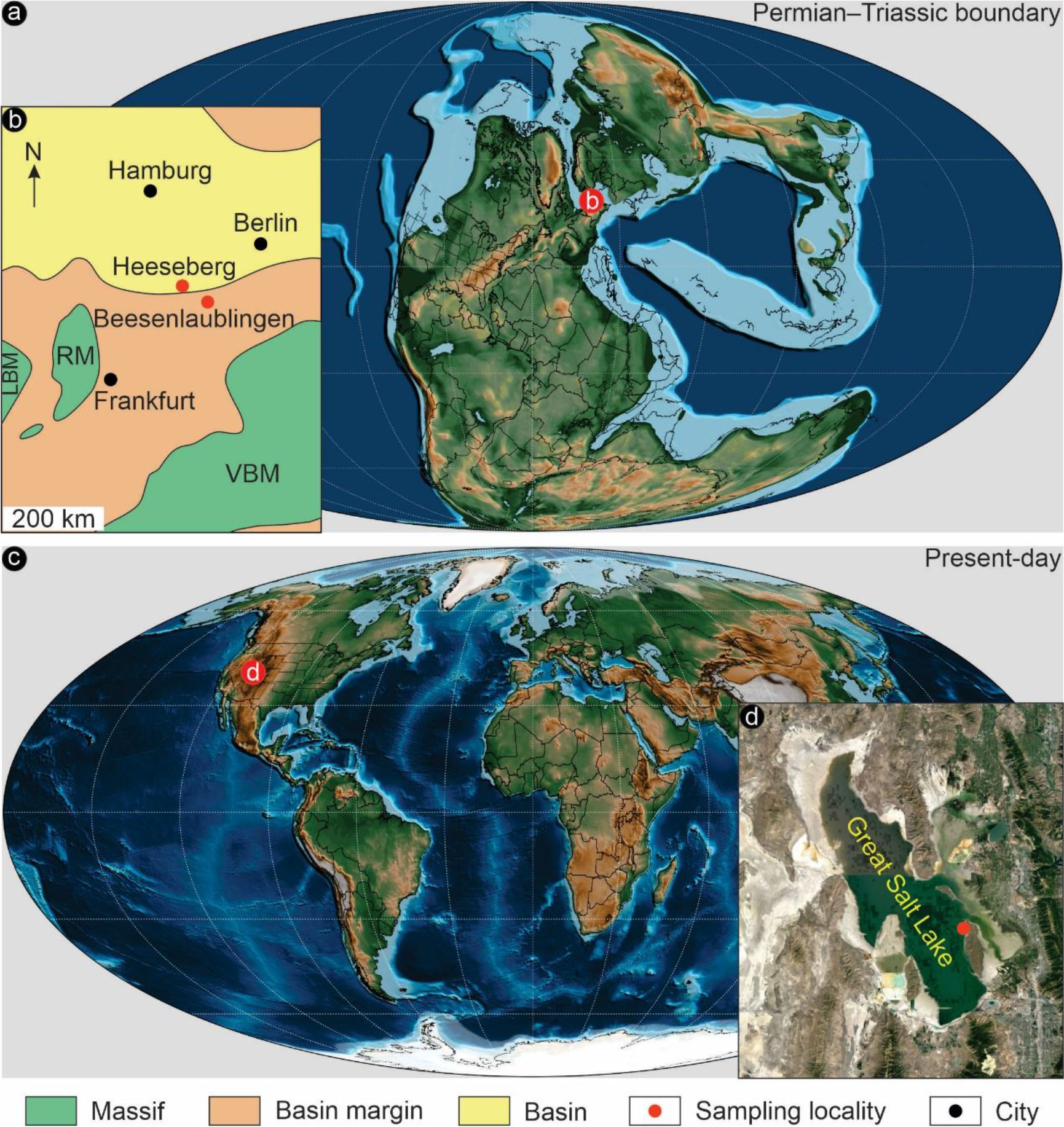
(a) Palaeogeographic map of Permian–Triassic boundary (∼250 Ma ago, Scotese, 2021). (b) Palaeogeography of the Lower Buntsandstein Subgroup (Early Triassic) in Germany (modified from Scholze et al., 2017) with main modern cities marked in black and the two sampling localities in red. LBM = London – Brabant – Massif. RM = Rhenish Massif. VBM = Vindelician – Bohemian – Massif. (c) Geographic map of Present-day (Scotese, 2021). (d) Geographic position of the sampling locality in red at the northwestern coast of Antelope Island in the Great Salt Lake (GSL) (Google Earth Pro image).

The GSL (Utah, USA) is an endorheic lake (Figure 1c) and a remnant of the Pleistocene Lake Bonneville (Oviatt et al., 1999; Shroder et al., 2016). Its salinity varies from approximately 12 % to 26 %, with Na^+^ and Cl^-^ being the major ions. The studied modern ooids were sampled at the northwestern coast of Antelope Island in 1994 and 1996 (Figure 1d), where the salinity is 12 % to 14 % (Chidsey et al., 2015). The samples are now stored in the Göttingen Geoscience Collections at the University of Göttingen.

### 2.2 Sample preparation

Petrographic thin sections were prepared from the Triassic samples of the GB. Modern ooid samples from GSL were processed directly after sampling in the field. Some samples were fixed in buffered formalin, dehydrated and stored in 70% ethanol, whole others were fixed in buffered glutardialdehyde (cooled on ice for 24 h) and postfixed with 2% osmium tetroxide (for details see Reitner, 1993). All GSL samples were then stained with Ca^2+^-chelating fluorescent dye (e.g., calcein) and non-fluorescent dye (e.g., alcian blue). Subsequently, thin sections were prepared from LR White-embedded specimens with a Leica SP 1600 saw microtome.

### 2.3 Petrography and analytical imaging

Thin sections were analysed using a Zeiss SteREO Discovery.V12 stereomicroscope and a Zeiss AXIO Imager. Z1 microscope. Photos were taken with an AxioCamMRc 5 MB camera.

For epifluorescence microscopy, a Zeiss AXIO Imager. Z1 microscope was utilized. It is equipped with a high-pressure mercury arc lamp (HBO 50, Zeiss; controlled by an EBX 75 ISOLATED electronic transformer) and a 10 AF488 filter (excitation wavelength = BP 450– 490 nm, emission wavelength = BP 515–565 nm). Thin sections with Ca^2+^-chelating fluorescent dye were studied with a ZEISS Axioplan using a high performance wide-band pass filter (blue, BP 450–490, LP 520; no. 487709) (for details see Hicks & Matthaei, 1958; Reitner, 1993).

For field emission scanning electron microscopy (FE-SEM), a Carl Zeiss LEO 1530 Gemini system was used. Some thin sections of the modern samples were etched by submerging them in a 5% ethylenediaminetetraacetic acid (EDTA) solution for 10–30 seconds.

A Bruker M4 Tornado instrument equipped with an XFlash 430 Silicon Drift Detector was used for micro X-ray fluorescence (μ-XRF) to obtain element distribution images of thin sections. Measurements (spatial resolution = 25 μm, pixel time = 8 ms) were conducted at 50 kV and 400 μA with a chamber pressure of 20 mbar.

A WITec alpha300 R fibre-coupled ultra-high throughput spectrometer was employed to collect Raman single spectra and spectral images to analyse mineral compositions of thin sections. Before analysis, the system was calibrated employing an integrated light source. For both single spectra and spectral images, the experimental setup includes a 532 nm excitation laser, an automatically controlled laser power of 20 mW, a 100×long working distance objective with a numerical aperture of 0.75, and a 300 g mm^-1^ grating. The spectrometer was centred at 2220 cm^-1^, covering a spectral range from 68 to 3914 cm^-1^. This setup has a spectral resolution of 2.2 cm^-1^. For single spectra, each was collected by two accumulations, with an acquisition time of 2 s. For spectral images, spectra were collected at a step size of 1 μm in horizontal and vertical direction by an acquisition time of 0.25 s for each spectrum. Automated cosmic ray correction, background subtraction and fitting using a Lorentz function were performed using the WITec Project software. Raman images were additionally processed with spectral averaging/smoothing and component analysis.

## 3 Results

### 3.1 Lower Triassic ooids from the GB

In the working area, the Bernburg Formation consists of siliciclastic and carbonate rocks which show abundant sedimentary structures (e.g., cross bedding, climbing ripples, wave ripples). A prominent feature of the formation are stromatolites with columnar growth forms and thicknesses of 10–50 cm (rarely up to 1.2 m) (Figure 2a,b). Ooids in the Bernburg Formation are typically large with diameters exceeding 2 mm (Figure 2c,d; see below for details). In many oolitic beds, ooids are overgrown by thin, 5–50 cm wide microbial crusts. Notably, these crusts show a lamination equivalent to that of the ooids. They either develop into a thicker stromatolite beds or are overlaid by another oolitic bed (Figure 2c,d).

**Figure 2.**
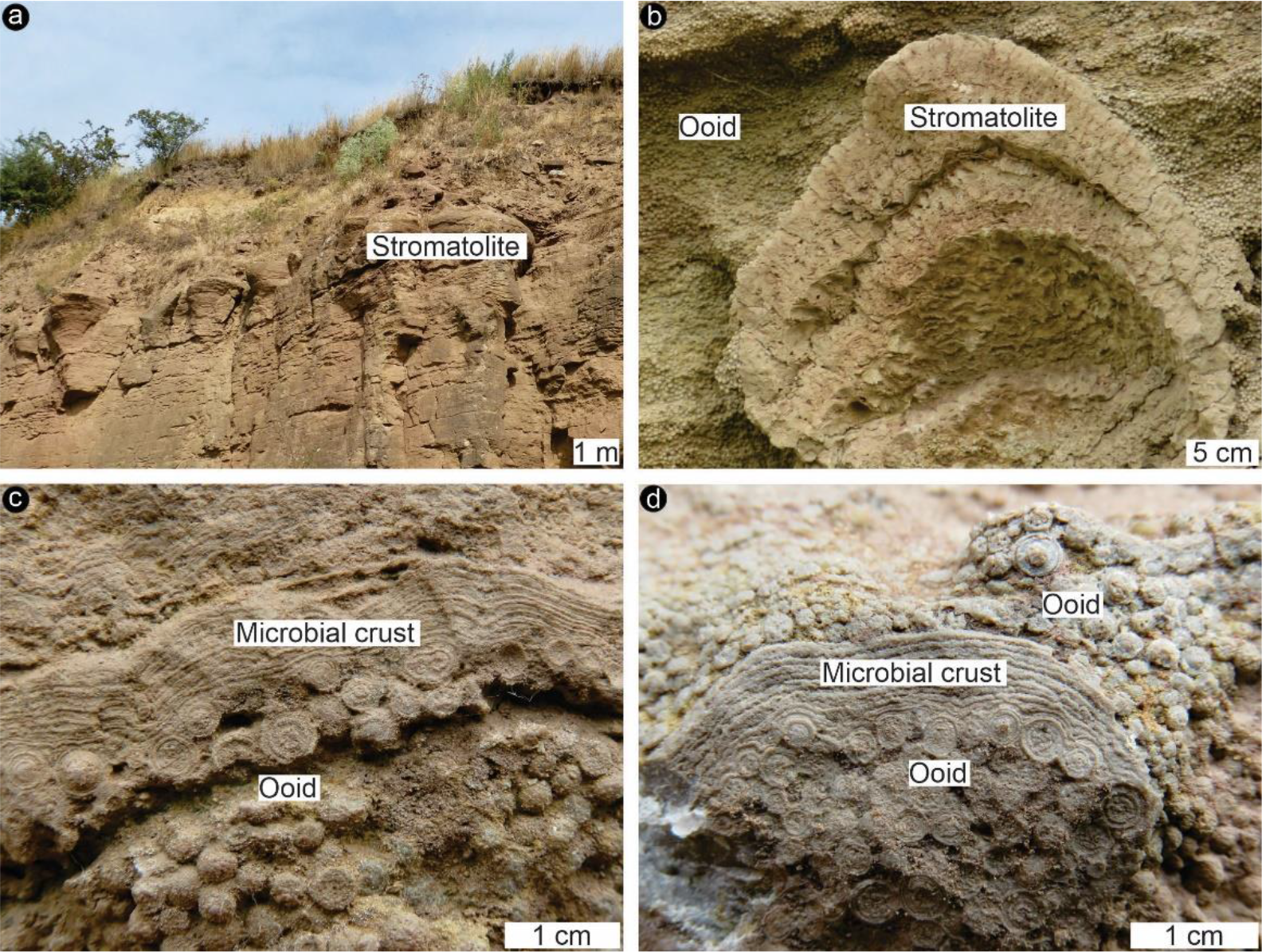
Field pictures of the Early Triassic, the Germanic Basin (GB). (a, b) Oolitic deposits alternate with columnar stromatolites. (c, d) The giant ooids, diameter > 2 mm, are overlaid by microbial crusts.

#### 3.1.1 Morphology and composition

The GB ooids are up to 4 mm in diameter and spherical to subspherical in shape. Their nuclei are most commonly dark micritic particles (e.g., Figure 3a) even though detrital (e.g., quartz) grains of various sizes are abundant in the sediment (e.g., Figures 4a, 5, 6). Four types of ooids are distinguished, according to the structure of their cortices. Type A ooids show co-occurring radial and tangential structures across their cortices (Figure 3a). Type B ooids exhibit alternating radial and tangential features (Figure 3b). Type C ooids are radial showing indistinct laminae (Figure 3c). Type D ooids are tangential exhibiting indistinct radial features (Figure 3d). Among the four types, Type A ooids are the most prominent. Type C ooids are typically smaller in size (< 0.5 mm) and can occur as the initial stage of other types of ooids. In some cases, adjacent ooids have coalesced during growth, forming “compound ooids” (Figure 3e-f). “Compound ooids” are very common in some samples.

**Figure 3.**
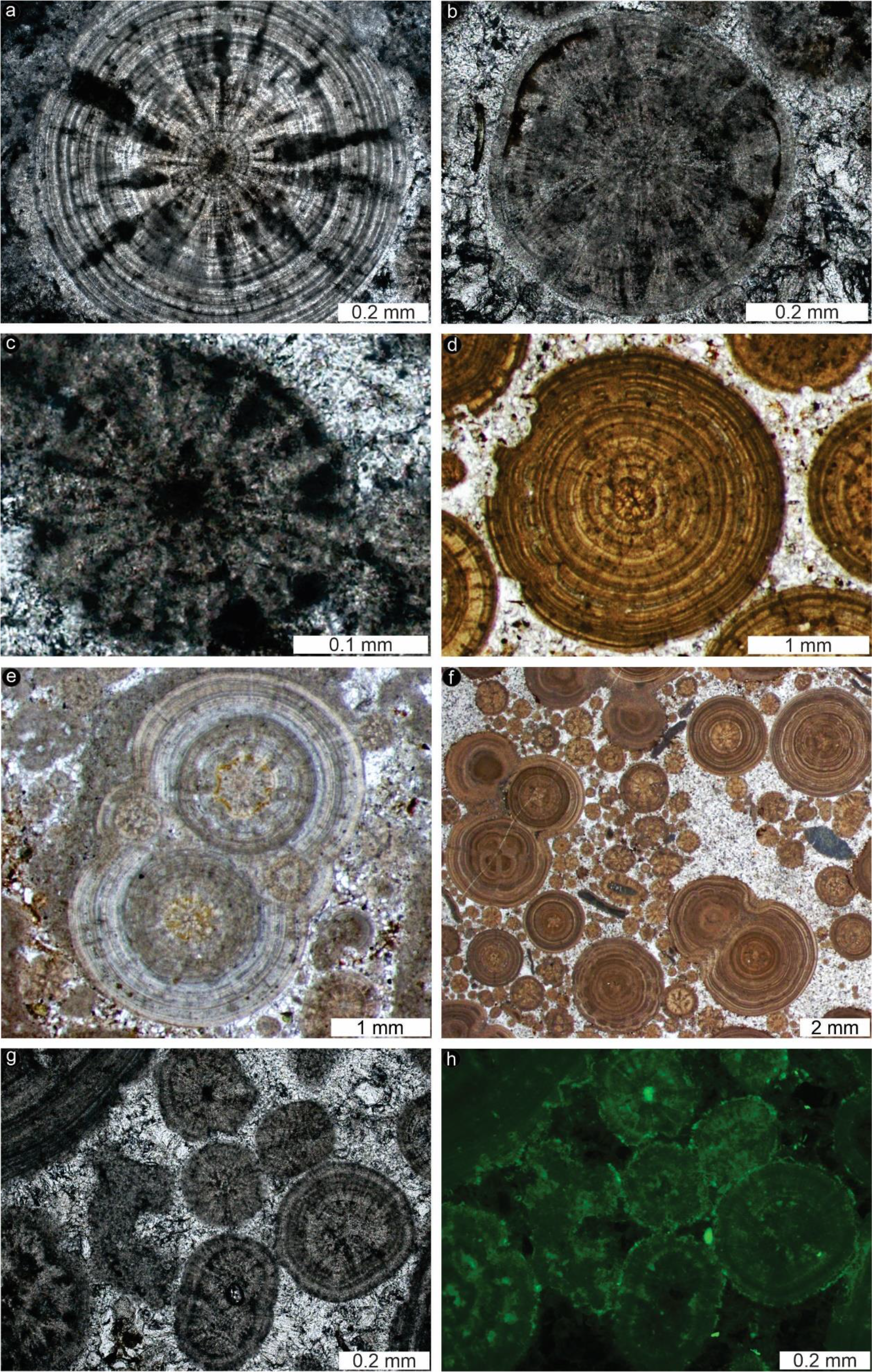
Thin section images of the Early Triassic ooids from GB. (a) Type A, characterized by co-occurring radial and tangential structures across their cortices. (b) Type B, with alternating radial and tangential features. (c) Type C, radial ooids showing indistinct laminae. (d) Type D, tangential ooids exhibiting indistinct radial features. (e-f) Compound ooids, formed by several ooids that have coalesced with each other during growth. (g-h) Same image under transmitted light (g) and fluorescence (h). Note that both radial and tangential cortices exhibit a strong green fluorescence, which is typically brighter in the darker radial segments and darker laminae.

**Figure 4.**
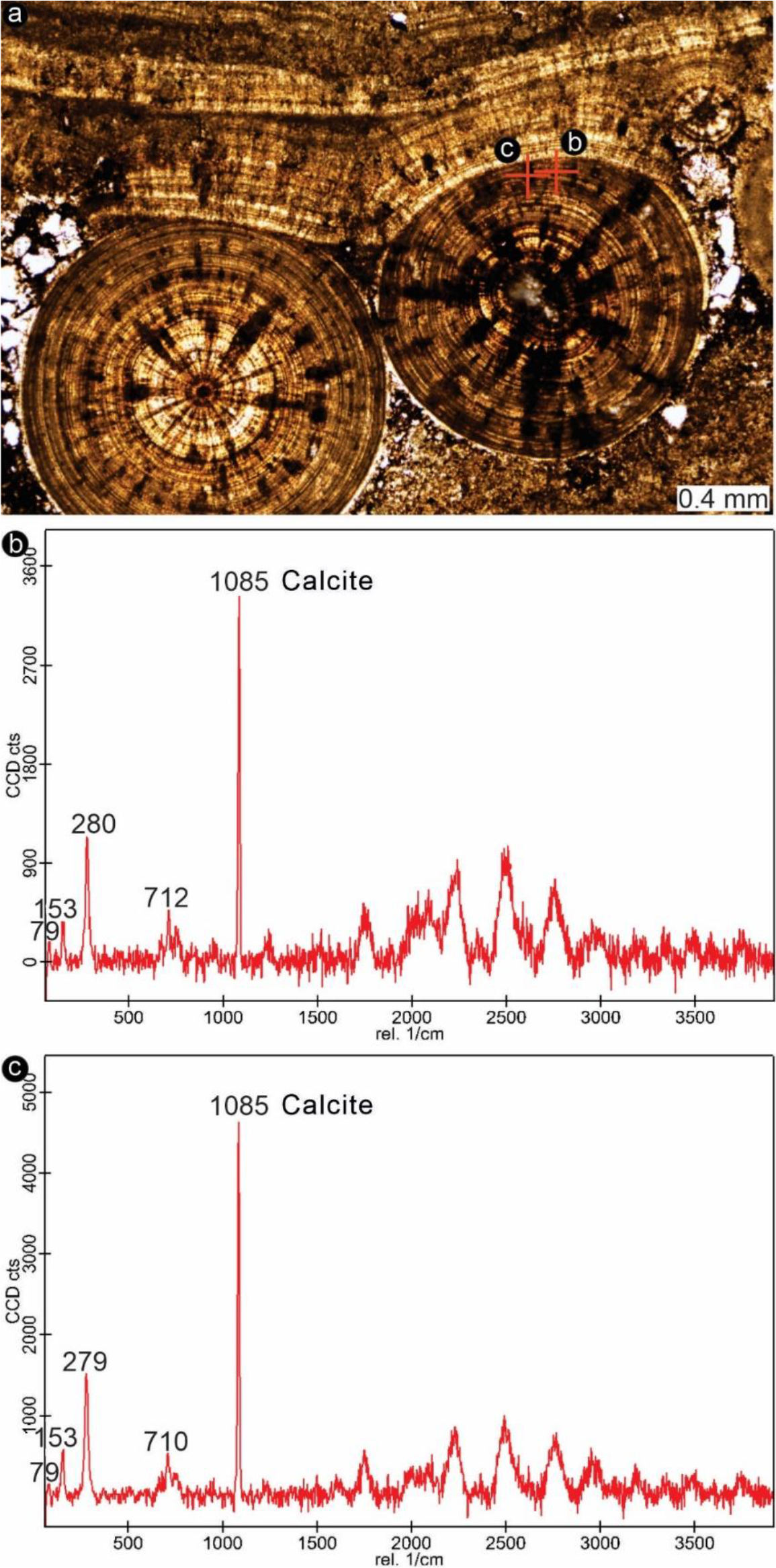
Raman spectroscopy (single spectra) images of the Triassic ooids from GB, showing that they are mainly composed of calcite. The unmarked peaks in (b, c) are attributed to fluorescence interference.

**Figure 5.**
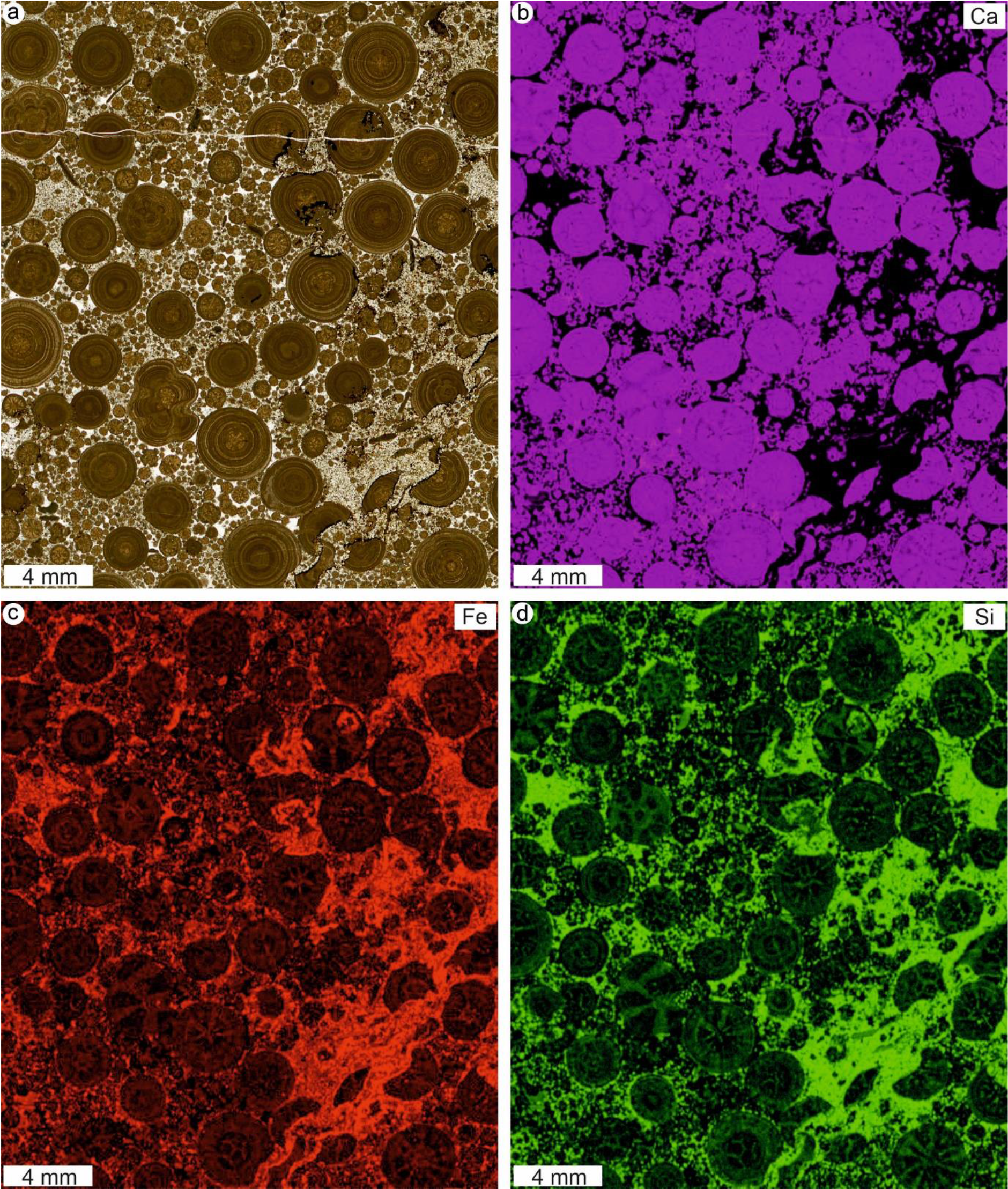
Micro X-ray fluorescence (μ-XRF) images of the Triassic ooids from GB. (a) Scan image (transmitted light). (b) Calcium (Ca) distribution. (c) Iron (Fe) distribution. (d) Silicon (Si) distribution

Both radial and tangential cortices exhibit a relatively strong green fluorescence, which is typically brighter in the darker radial segments and darker laminae (Figure 3g,h). Raman spectroscopy shows that the ooids are mainly composed of calcite (Figure 4). The dark and light radial segments display different contents of elements, e.g., Calcium (Ca) (Figure 5a,b), Iron (Fe) (Figure 5a,c) and Silicon (Si) (Figure 5a,d), as indicated by μ-XRF. The matrix between ooids is typically enriched in Si, which is due to the presence of abundant quartz grains. At the same time, there is no petrographic or μ-XRF evidence for the presence of quartz grains in the center of ooids.

#### 3.1.2 Relationship between ooids and microbial crusts/stromatolites

The studied GB ooids are often overlaid by laterally-continuous microbial crusts equivalent to the initial stage of stromatolites (Figure 2c,d). Ooids and microbial crusts also alternate on a mm-scale (Figure 6a). In other cases, microbial crusts completely envelope clusters of ooids (Figure 6b), which was already described by Kalkowsky (1908) as “ooid bags” (Ooid Beutel) and also observed by other researchers (Paul & Peryt, 2000; Paul et al., 2011).

**Figure 6.**
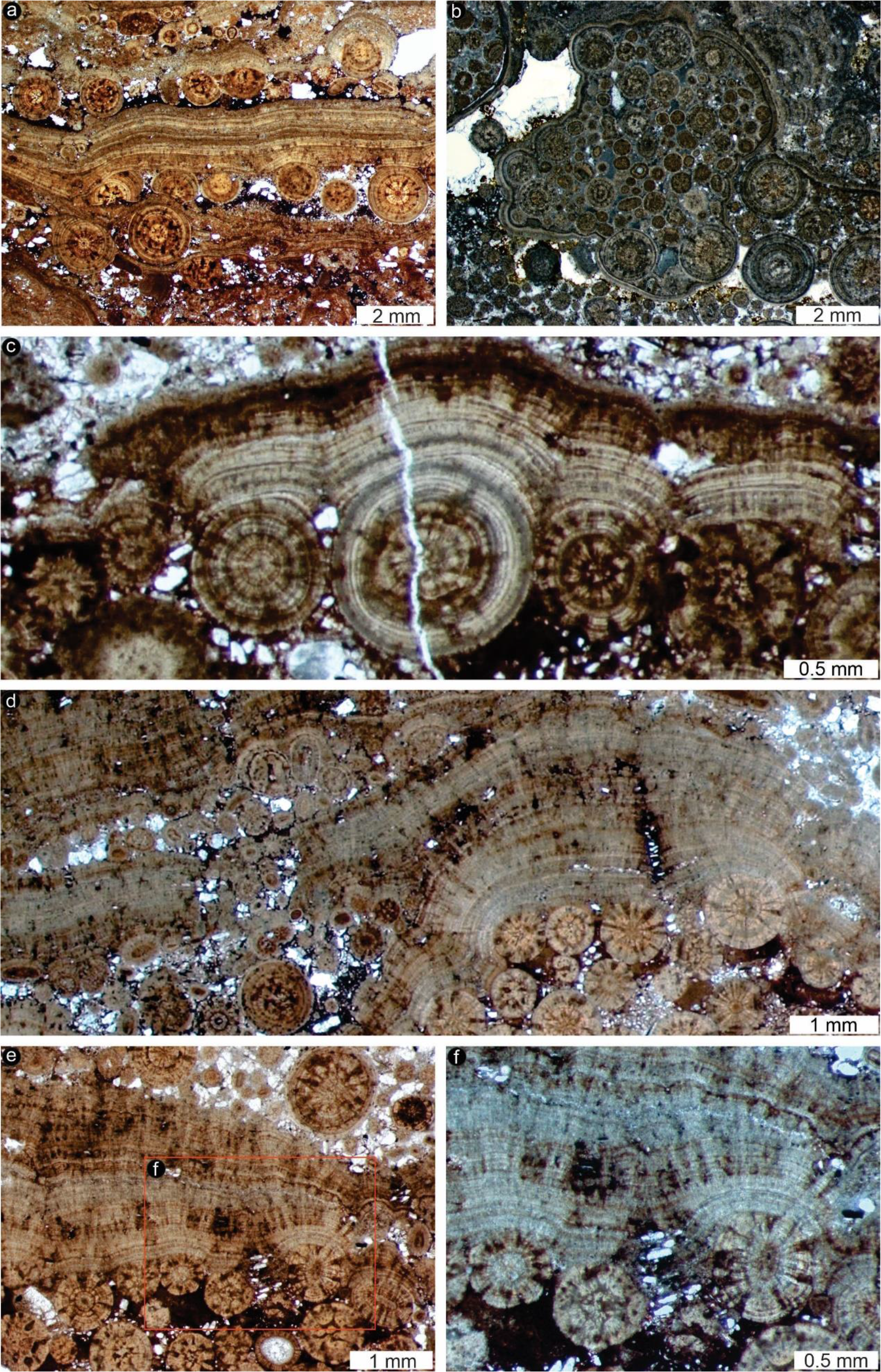
Thin section images of the ooids and microbial crusts/stromatolites of the Early Triassic from GB. (a) Ooids and microbial crusts alternate on a mm-scale. Truncation is observed between the ooid cortices and the overlying microbial crusts. (b) Microbial crusts completely envelope clusters of ooids. (c, d) The microfabric of both ooid laminae and stromatolite laminae is almost identical, with laminae showing a characteristical internal palisade structure composed by thin crystals growing perpendicularly to lamination. (e, f) The microbial crusts also show structures equivalent to the radial and tangential structures of ooid cortices. The rectangle in (e) is magnified as (f).

Furthermore, the microfabric of both ooid laminae and stromatolite laminae is almost identical, with laminae showing a characteristical internal palisade structure composed by thin crystals growing perpendicularly to lamination (Figure 6c,d). In addition, the microbial crusts also show structures equivalent to the radial and tangential structures of ooid cortices (Figure 6e,f). Locally, truncation is observed between the ooid cortices and the overlying microbial crusts, but the microfabric of laminae remains equivalent (Figure 6a).

### 3.2 Modern ooids from the GSL

#### 3.2.1 Morphology and composition

GSL deposits contain abundant ooids and associated microbial crusts (Chidsey et al., 2015; Vennin et al., 2019) (Figure 7a). The studied GSL ooids range from 0.2 mm to 1 mm in diameter and are ellipsoidal to subspherical in shape (Figures 7-9). The nuclei of ooids are either micritic particles or detrital (quartz or feldspar) grains (Figures 7,8c and 9a). Ooid cortices are aragonitic, as indicated by Raman spectroscopy (Figure 8 d-f).

**Figure 7.**
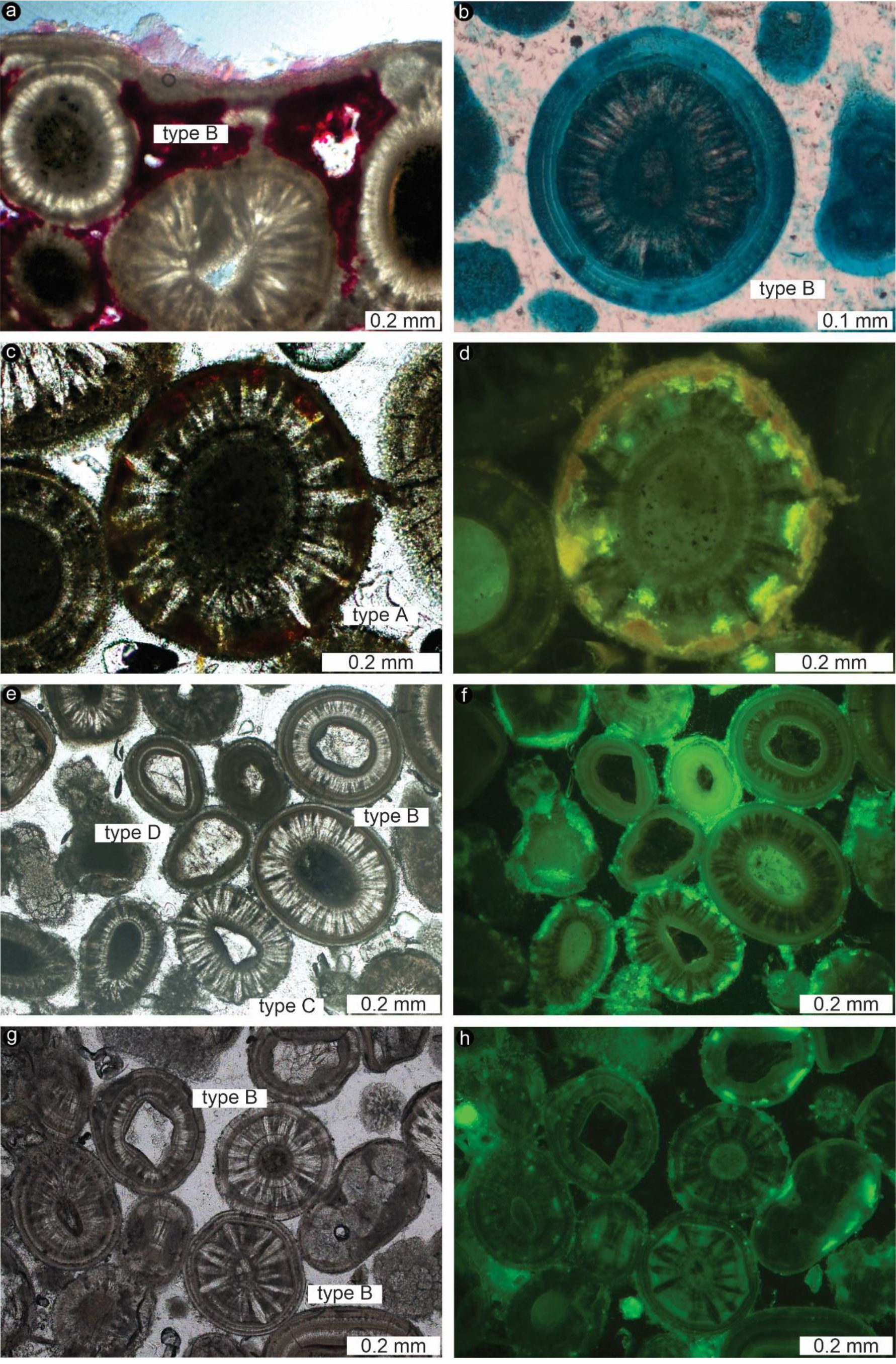
Thin section images of the ooids and/or microbial crusts from GSL. Note that their cortex structures are similar to those of the Triassic specimens. (a) Ooids, with cortices showing alternating radial and tangential features (Type B), which are overlaid by thin microbial crusts (basic fuchsin stain). (b) Type B ooid, with alternating radial and tangential features. Note that alcian blue staining is stronger in tangential cortices than that in radial ones. (c) Type A ooid, characterized by co-occurring radial and tangential structures across their cortices. (d) Same image as (c), under fluorescence. Note that, from the outer edge to the inside, orange, yellow and green colors are observed, suggesting declining abundance of free Ca^2+^ and acidic OM, corresponding with a gradual increase in mineralization. (e, g) Ooids corresponding to Type B (alternating radial and tangential features in the cortex), C (radial ooids showing indistinct laminae) and D (tangential ooids exhibiting indistinct radial features). (f, h) Same images as (e, g), under fluorescence. Significant internal differences in the intensity of fluorescence are shown. Tangential cortices have a brighter green fluorescence than those with radial structures. In the radial ones, the dark radial segments show a brighter green fluorescence than the lighter segments. Occasionally, a very bright green rim is observed at the outer edge.

**Figure 8.**
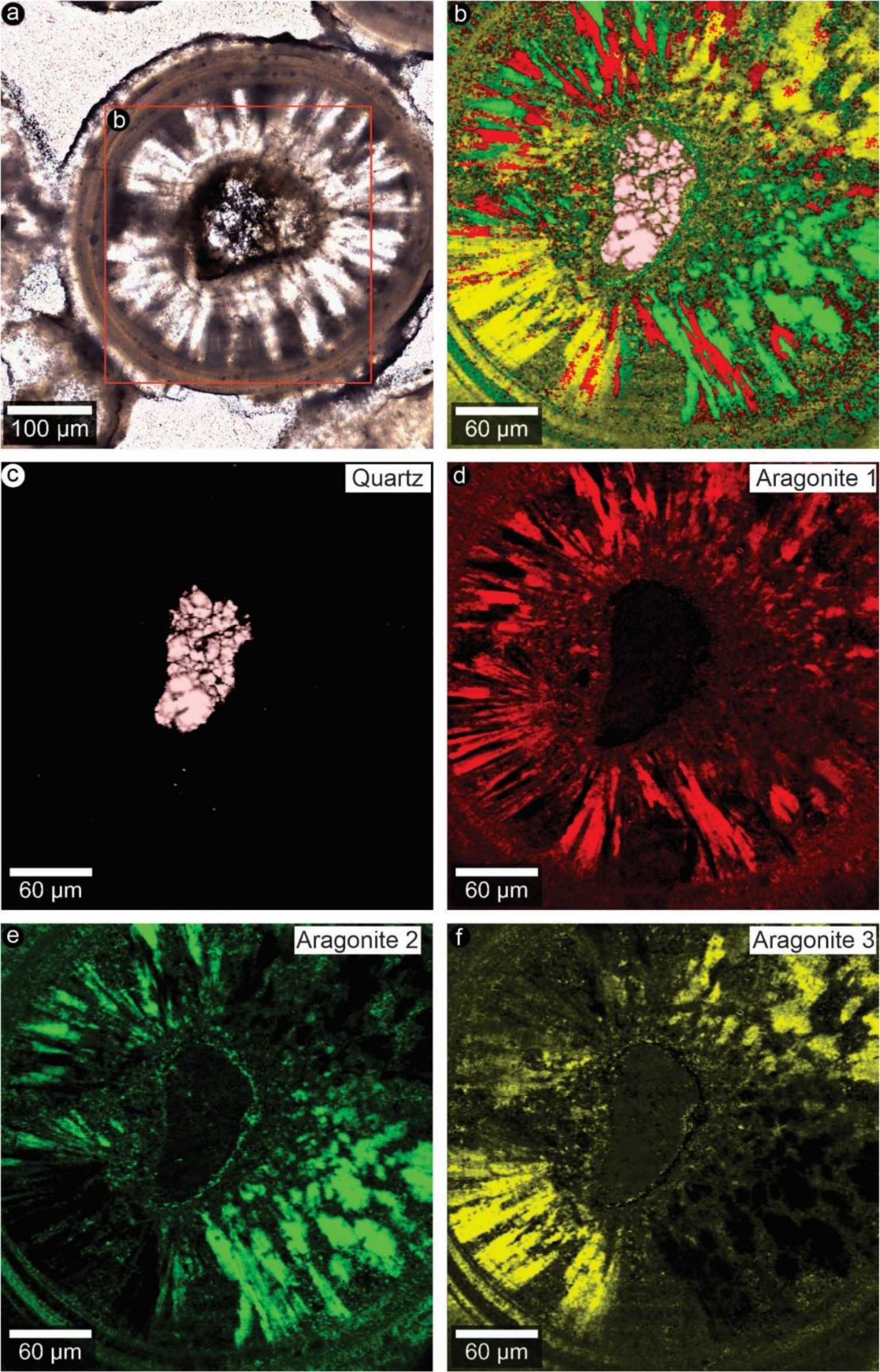
Raman spectral images of modern ooids from GSL. (a) Thin section image (transmitted light). (b) Combined image. (c) Quartz as the core. (d-f) Different preferred orientations of aragonite crystals. The rectangle in (a) is magnified as (b).

**Figure 9.**
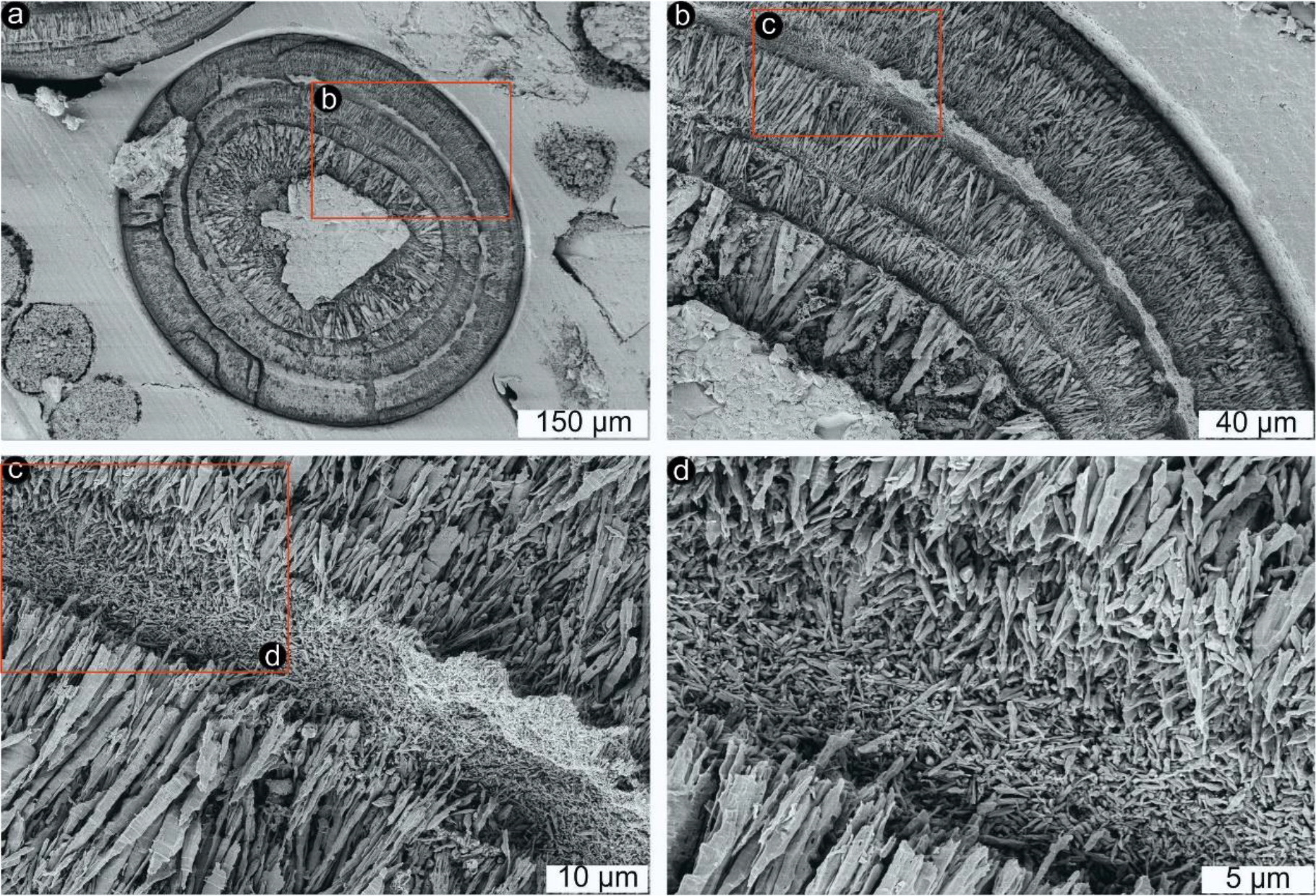
FE-SEM images of modern ooids from GSL. (a-d) Radial parts of the ooid cortices consist of > 50 µm long fan-shaped crystals. Tangential parts are formed by alternating laminae of radial crystals and smaller (< 5 µm) needles with irregular arrangement. The rectangle in (a) is magnified as (b). The rectangle in (b) is magnified as (c). The rectangle in (c) is magnified as (d).

GSL ooids can be classified based on their cortex structures and include the same types as the Triassic deposits from the GB (Figure 7a-c,e,g). Type A ooids show co-occurring radial and tangential structures across their cortices (Figure 7c). Type B ooids exhibit alternating radial and tangential features (Figure 7a,b,e,g). Type C ooids are radial showing indistinct laminae (Figure 7e). Type D ooids are tangential exhibiting indistinct radial features (Figure 7e). Raman spectroscopy indicates different preferred orientations of aragonite crystals in radial and tangential parts of the cortices (Figure 8d-f). FE-SEM shows that radial parts of the cortices consist of > 50 µm long fan-shaped crystals. Tangential parts are formed by alternating laminae of radial crystals and smaller (< 5 µm) needles with irregular arrangement (Figure 9). “Compound ooids” are rarely observed.

#### 3.2.2 OM in ooids

GSL ooids show significant internal differences in the intensity of green fluorescence (Figure 7e-h). Tangential cortices have a brighter green fluorescence than those with radial structures, and in the radial ones, the dark radial segments show a brighter green fluorescence than the lighter segments (Figure 7e-h). Occasionally, a very bright green rim is observed at the outer edge (Figure 7e,f). Alcian blue staining, which indicates the presence of carboxylic groups in acidic polysaccharides, is stronger in tangential cortices than in radial ones (Figure 7b).

Fluorescent dye calcein detects free Ca^2+^ in acidic OM, and the concentrations of free Ca^2+^ are reflected by fluorescence intensities with different colors (Reitner et al., 1997). In the GSL ooids, from their outer edge to their inside, orange, yellow and green colors are observed (Figure 7c,d), suggesting declining abundance of free Ca^2+^ and acidic OM, corresponding with a gradual increase in mineralization.

## 4 Discussion

### 4.1 Comparison of the GB and GSL ooids

The GB and GSL ooids have some obvious differences, mainly size (GB ooids are typically much larger than GSL ooids) and shape (GB ooids are mainly spherical while GSL ooids are predominantly ellipsoidal) (Figures 3 and 7). Furthermore, GB ooids rarely have detrital grains as nuclei, while this is not the case in GSL ooids (Figures 3-9). This is notable since both deposits are rich in detrital (mainly quartz) grains.

At the same time, GB and GSL ooids share remarkable similarities, which is why they have been previously compared (Käsbohrer & Kuss, 2019; Pei 2022). For instance, GB and GSL ooids display the same types of cortices, that is, cortices with radial structure (Type C), cortices with tangential structure (Type D) and cortices with either a mixture or an alternation of both structures (Type A and B, respectively) (Figures 3 and 7). Besides, ooids from both settings have micritic particles as nuclei and are rich in radial features (Figures 3 and 7). Fluorescence microscopy indicates the presence of OM in ooids from the GB (Figure 3g,h) and the GSL (Figure 7). However, OM contents seem to be much lower in case of GB ooids, which is likely due to degradation through geological time.

Ooid types similar to those from the GB and GSL have been observed in modern hypersaline environments (Friedmann et al., 1973, 1985; Krumbein & Cohen, 1974; Krumbein, 1983; Gerdes et al., 1994, 2000; Hubert et al., 2018; Loreau & Purser, 1973; Strasser, 1986; Suarez-Gonzalez & Reitner, 2021). In an agreement with the geological settings and water chemistry, ooids from both deposits investigated herein are supposed to have formed under hypersaline conditions as well (Paul & Peryt, 2000; Chidsey et al., 2015; Käsbohrer & Kuss, 2019).

### 4.2 Formation of ooids statically versus dynamically

The presence of abundant sedimentary structures (e.g., cross bedding, climbing ripples and wave ripples) and abundant detrital (mainly quartz) grains in the GB and GSL deposits indicate reworking and transportation processes (e.g., Reitner et al., 1997; Paul & Peryt, 2000; Käsbohrer & Kuss, 2019) (Figures 4 and 6). Yet, various lines of evidence suggest that ooids from both deposits were not formed dynamically during transportation. For instance, quartz grains are abundant constituents of the deposits, but almost never or partially form the nuclei of the ooids (Figures 3-9). Instead, the nuclei of ooids typically consist of small micritic particles, which are themselves rare constituents of the oolitic deposits. Furthermore, the GB deposits contain abundant “compound ooids” consisting of ooids that coalesced with each other during growth, which is inconsistent with growth through constant motion as bed-load or in suspension. These features (micritic nuclei in detrital matrix and cortex coalescence) have recently been shown as characteristic of modern ooids growing *in situ* within microbial mats (Suarez-Gonzalez & Reitner, 2021). At the same time, ooids and microbial crusts are closely related and their microstructures are commonly identical, indicating that similar processes were involved in their formation (e.g., Figures 6c-f and 7a). Taken together, the observed features suggest that these ooids develop statically (perhaps even within microbial mats) (e.g., O’Reilly et al., 2017; Mariotti et al., 2018; Anderson et al., 2020), rather than form around transported grains in agitated environments (e.g., Duguid et al., 2010; Trower et al., 2018).

### 4.3 Organic influence on ooid formation

Abundant OM within GSL ooids (Figure 7b,e-h), as well as increasing OM contents associated with decreasing mineralization degrees from the nuclei towards to the outer edge of ooids (Figure 7c,d), may indicate a relationship between OM and carbonate formation. In case of GB ooids, the possible relationship between OM and carbonate minerals is less obvious, likely because OM has been degraded. However, as argued before, microbial crusts in the GB deposit typically represent a continuation of the microfabric of the ooid cortices (e.g., Figure 6c-f), suggesting similar formation processes. Notably, Kalkowsky (1908), in the same ground- breaking study where he coined the term “stromatolite”, already used this to argue for an organic influence on the origin of ooids in the GB, stating that “*if somebody insists on the inorganic genesis of ooids, he will, however, never find a good explanation for* [the fact that] (…) *ooids often form the initiating stage of stromatolites*” (translated by Krumbein, 1983; Paul et al., 2011), which underlines the great value and continuing impact of this observation-based research. In addition, Granier & Lapointe, (2021, 2022) recently argued a microbial influence on the development of other examples of fossil ooids (very similar to the Triassic examples studied herein). Actually, the organic origin of these crusts with fibrous-palisade microstructure is hard to establish, and they have been previously interpreted as mostly inorganic, although some microbial influence was not ruled out (Paul & Peryt, 2000). It is true that in modern environments, similar fibrous microfabrics and crusts occur during moments of environmental increase in CaCO3 supersaturation, but typically within microbial communities (e.g., Suarez-Gonzalez & Reitner, 2021; Shen et al., 2022), and they have also been observed within fossil microbialites (Camoin et al., 1997; Kirkham & Tucker, 2018).

### 4.4 Possible ooid formation processes

Various factors have been traditionally proposed for explaining differences in the cortex structures of ooids, mainly dealing with different environmental conditions (e.g., Strasser, 1986). Given the organic influence on ooid formation highlighted herein, other hypotheses related with organic matter can be proposed. One important function of microbial EPS is the inhibition of mineral precipitation (e.g., Decho, 2010). In addition, organic matter can function as nucleation site for mineralization (“Organomineralization”: Mitterer, 1968; Suess & Fütterer, 1972; Ferguson et al., 1978; Trichet & Défarge, 1995; Reitner et al., 1995a, b, 1997; Reitner, 2004). The observed different intensities of alcian blue staining (indicative for the presence of carboxylic groups (COO^-^) in acidic polysaccharides) and green fluorescence (likely indicative for the content of OM) between tangential and radial cortices from GSL (Figure 7b, f, h) suggest potential differences of OM in forming both cortices. Potentially degraded organic matter is more involved in forming radial cortices while fresher EPS is more related with tangential cortices (Figure 10). However, further research on OM associated with different types of ooids is needed.

**Figure 10.**
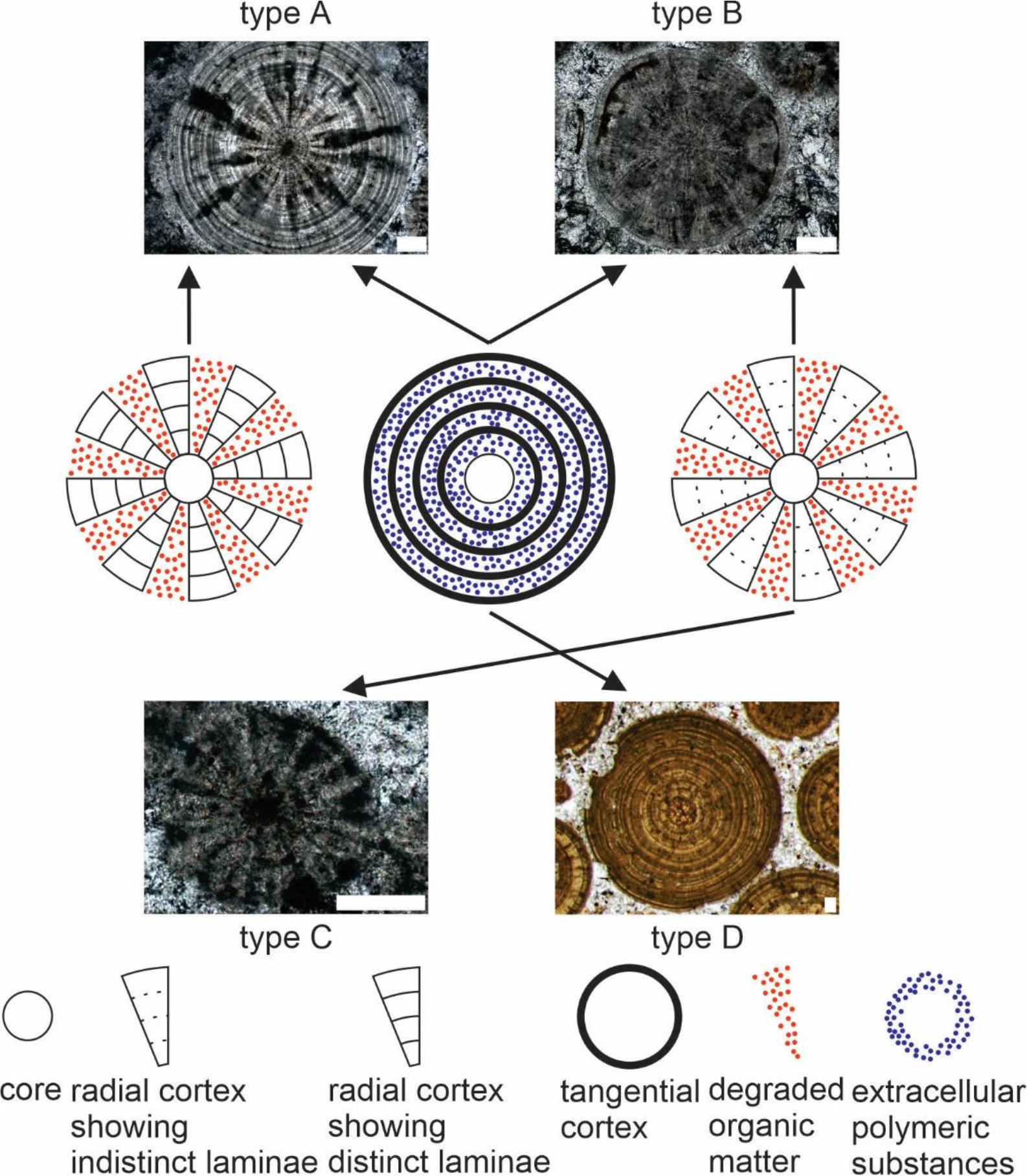
Extracellular polymeric substances (EPS) and degraded organic matter (OM) shape four types of ooids. Potentially degraded organic matter is more involved in forming radial cortices while fresher EPS is more related with tangential cortices. All scale bars represent 0.1 mm.

## 5 Conclusion

We studied ooids in deposits from the GSL and GB, two world-famous geobiological sites. Ooids in both localities display extreme similarities in their internal microstructures, suggesting that they could have been formed by similar processes. The micritic ooid nuclei, the common presence of “compound ooids” as well as identical characteristics of ooid laminae and overlaid microbial crusts are all supportive of static growth. OM is present in interspaces between radial and tangential ooid cortices and becomes more abundant towards the outer edge in case of radial ooids. Together, these features strongly indicate that acidic OM played a key-role in the formation of ooids. This is additionally supported by the close association of ooids and microbial crusts in the GB deposit, since the latter typically form through mineral precipitation associated with OM.

## Acknowledgements

Axel Hackmann and Wolfgang Dröse are thanked for lab assistance. This study is financially supported by the China Council Scholarship (CSC), a Teach@Tübingen Fellowship from the University of Tübingen and Göttingen Academy of Sciences and Humanities. PSG acknowledges funding by a postdoctoral “Humboldt Research Fellowship” of the Alexander von Humboldt Foundation.

## Notes

### Competing Interest Statement

The authors have declared no competing interest.

